# A new strategy for identifying polysialylated proteins reveals they are secreted from cancer cells as soluble proteins and as part of extracellular vesicles

**DOI:** 10.1101/2022.09.01.506237

**Authors:** Carmanah Hunter, Tahlia Derksen, Julieanna Karathra, Kristi Baker, Mark Nitz, Lisa M. Willis

**Affiliations:** Department of Biological Sciences, University of Alberta, Edmonton, AB, T6G2E9; Department of Experimental Oncology, University of Alberta, Edmonton, AB, T6G2E9; Department of Medical Microbiology and Immunology, University of Alberta, Edmonton, AB, T6G2E9; Department of Chemistry, University of Toronto, Toronto, ON, M5S 3H6

**Keywords:** ManNAz, metabolic engineering, polysialic acid, cancer, QSOX2, AGR2, GOLPH4

## Abstract

Polysialic acid (polySia) is a long homopolymer consisting of α2,8-linked sialic acid with tightly regulated expression in humans. In healthy adults, it occurs on cell surface glycoproteins in neuronal, reproductive, and immune tissues; however, it is aberrantly present in many cancers and its overexpression correlates with significantly increased metastasis and poor prognosis. Prompted by the observation that the MCF-7 breast cancer cell line contains only intracellular polySia, we investigated the secretion of polySia from MCF-7 cells. PolySia was found predominantly on soluble proteins in MCF-7 conditioned media, but also on extracellular vesicles (EVs), secreted from the cells. Since MCF-7 cells do not express known polysialylated proteins, we developed a robust method for purifying polysialylated proteins that uses a metabolic labelling strategy to introduce a bioorthogonal functionality into polySia. Using this method we identified three previously unknown polysialylated proteins, and found that two of these proteins - AGR2 and QSOX2 – were secreted from MCF-7 cells. We confirmed that QSOX2 found in EV-depleted MCF-7 cell conditioned media was polysialylated. Herein we report the secretion of polysialic acid on both soluble and EV-associated proteins from MCF-7 cancer cells and introduce a new method to efficiently identify polysialylated proteins. These findings have exciting implications for understanding the roles of polySia in cancer progression and metastasis and for identifying new cancer biomarkers.

## Introduction

Polysialic acid (polySia) is a linear polymer of α2,8-linked sialic acid residues that is of fundamental biological interest due to its pivotal roles in the regulation of the nervous, immune, and reproductive systems in healthy human adults. PolySia elaborates N- and O-linked glycans on a small subset of proteins and imparts profound consequences on the proteins and cells to which it is attached (1–3). Cells containing polysialylated proteins are typically more migratory, and polySia is required for axonal migration (4), neurite outgrowth (5), and dendritic cell migration (6). PolySia also acts as an immune checkpoint and attenuates the immune response, similar to what has been observed for other sialosides (7, 8). PolySia on macrophages decreases their phagocytic capacity (9) as well as the ability of dendritic cells to induce T cell proliferation and release proinflammatory cytokines (10). Exogenous polySia is also anti-inflammatory in a humanized mouse model of macular degeneration (11). However, the majority of details regarding the roles of polySia, particularly in the immune and reproductive systems, have not been elucidated.

In addition to its critical roles in healthy humans, polySia is dysregulated in several chronic diseases and contributes to their pathology. Emerging evidence suggests that polySia plays an etiology in some mental illnesses and neurodegenerative diseases. Single nucleotide polymorphisms (SNPs) occur in ST8SiaII, one of the two enzymes responsible for polySia biosynthesis, in patients with schizophrenia, autism spectrum disorders and bipolar disorder (12–15). Aberrant levels of polySia-NCAM in brain tissue, cerebrospinal fluid, and serum have been found in those with Alzheimer’s disease, Parkinson’s disease, depression, and schizophrenia (16–21). Furthermore, the link between polySia and metastasis, the cause of the vast majority of cancer mortalities, is well established though the mechanisms have yet to be elucidated (22–25). High expression levels of polySia in clinical samples (tumor tissue sections) correlates with significantly increased metastasis and poorer prognoses (22, 23). *In vitro* models have demonstrated that polySia is required for cell migration in the neuroblastoma SH-SY5Y and pancreatic carcinoma PANC-1 cell lines (26–29), and the requirement of polySia for cell migration and metastasis has been confirmed *in vivo* (24, 28, 30, 31). Given the extensive evidence of a role for polySia in cancer progression, it is imperative to understand its mechanisms of action.

Cancer cells secrete extracellular vesicles (EVs) and soluble components which together comprise the cancer secretome (32). The cancer secretome has been identified as critical for multiple aspects of cancer progression, including immune evasion (33, 34), extracellular matrix remodeling (35, 36), and formation of the pre-metastatic niche (37, 38). Study of the cancer secretome not only has the potential to generate non-invasive diagnostic biomarkers for cancers, but could also provide insight into cancer progression (39). PolySia was recently identified in intracellular pools that are released from healthy cells (40, 41). Werneburg et al found that intracellular pools of Golgi-associated polySia could be depleted from microglia and oligodendrocyte precursor cells and increased in cell culture supernatant upon cell activation (41). Strubl et al demonstrated that intracellular pools of polySia in human umbilical vein endothelial cells (HUVECs) could be secreted into media (40). Additionally, polySia is present in serum (42) where it has been identified on EVs (43). We hypothesized that polysialylated proteins are also components of the cancer secretome and may represent a novel population of polysialylated proteins.

Investigating polySia can be a challenge. While there are excellent α-polySia antibodies available, specialized conditions are required for immunoblotting free polySia (44). For analysis of polysialylated proteins, the large, anionic nature of the polymer frequently interferes with protein mobility and protein detection via immunoblotting (41), making it difficult to demonstrate that a protein is polysialylated. Indeed, only neural cell adhesion molecule (NCAM/CD56), synaptic cell adhesion molecule (SynCAM), neuropilin-2, and the polysialyltransferases ST8Sia2 and ST8Sia4 have unequivocal evidence for their polysialylation (3). Proteins which require additional evidence for their polysialylation or for which polysialylation is controversial are E-selectin ligand 1 (41), CCR7 (6, 45), CD36 (46), and the voltage gated sodium channel (47). PolySia length can be distinguished through fluorescent labeling with the DMB and separation by reverse phase chromatography (48). However, the labeling is performed under acidic conditions potentially affecting the length and amount of polySia as the glycan is acid-labile (49). A recent sandwich ELISA has been described that allows for the quantification of polySia as well as specific polysialylated proteins under physiological conditions (42). This polySia ELISA has the benefit of being fast, specific to a polysialylated protein, tolerant of complex mixtures such as serum samples, and amenable to low abundance polysialylated proteins. The same properties that make polySia difficult to detect also make identification of polysialylated proteins via pull down approaches challenging. Many proteins, particularly basic proteins, tend to interact non-specifically with polySia and cannot be removed under the mild washing conditions of immunoprecipitation. The resulting proteomics experiments often contain numerous hits which require either a substantial number of antibodies for validation or a biased selection of hits to validate based on known function and subcellular localization (10, 50). The reciprocal affinity purification has been used to validate polysialylated proteins previously but, like all polySia analysis techniques, suffers from its own drawbacks. For example, a protein with shorter/fewer polySia chains might not display the same kind of large molecular weight shift that is observed for heavily polysialylated proteins like NCAM. Even if a substantial shift were to occur on a given polysialylated protein, if only a fraction of said protein were polysialylated, the non-polysialylated fraction would confound any immunoblotting done in a reciprocal affinity purification analysis. Due to these issues, previous identification of polysialylated proteins has sometimes had to rely on manipulation of the system to demonstrate polysialylation, e.g. overexpression of the polysialyltransferase (45), which shows that a given protein *can* be polysialylated, but not necessarily that it *is* under normal conditions. Improved methods to identify polysialylated proteins would substantially enhance our ability to investigate the mechanisms underpinning the role of polySia in health and disease.

Here we demonstrate that polySia is indeed secreted from the MCF-7 breast cancer cell line as we detected polySia both as a component of EVs and on soluble proteins. We developed a robust method for purifying polysialylated proteins that uses a metabolic labelling strategy to introduce a bioorthogonal functionality into polySia. The biorthogonal handle allows robust washing of the isolated conjugates after selective elution from a lectin affinity column. Proteomics analysis of the isolated proteins allowed identification of three previously unknown polysialylated proteins in MCF-7 cells. These proteins were validated by polySia ELISA (42) and immunoblotting. We found that two of the proteins, QSOX2 and AGR2, are secreted from MCF-7 cells, and confirmed that secreted QSOX2 is polysialylated. These findings provide a starting point for better understanding the molecular roles of polySia in cancer biology.

## Materials and Methods

### Cell lines, serum and antibodies

MCF7 (ATCC HTB-22) were obtained from ATCC through Dr. Christina Allen. MCF-7 and were maintained in D-MEM supplemented with 10% FBS (Sigma), 100 U/mL penicillin, and 100 μg/mL streptomycin. Antibodies used in this study are α-polySia mAb 735 (BioAspect BA-Ab00240-2.0), α-NCAM (Santa Cruz sc-7326), α-QSOX2 (Abcam ab191168 and ab121376), α-GOLIM4 (Abcam ab181849), α-AGR2/3 (Santa Cruz sc-376653), α-G6PD (Santa Cruz sc-373886), α-MGAT2 (Abcam ab184965), α-LRP2 (Santa Cruz sc-515772), α-SDF4 (Santa Cruz sc-393930), GM130 (Cell Signaling 12480T), α-PDI (Cell Signaling 3501T), α-CD63 (abcam ab8219), α-mouse IgG-HRP (Cell Signaling 7076S), and α-rabbit IgG-HRP (Cell Signaling 7074S).

### Protein expression

GFP-EndoN_DM_ (51, 52) was expressed in BL21 (DE3). Overnight cultures were used at 1/100 dilution to inoculate LB media supplemented with 100 μg/mL ampicillin. Cultures were grown at 37°C for 2 h with shaking, then 20 °C for 30 min before inducing with IPTG to a final concentration of 0.5 mM. 26 h after addition of IPTG, cells were harvested by centrifugation, resuspended in 20 mM Tris HCl, 500 mM NaCl, 10 mM imidazole, 10% glycerol, 10 mM β-mercaptoethanol, pH 8 and lysed. Protease inhibitor cocktail (Roche) and Benzonase (Sigma) were added and the suspension was cleared by centrifugation at 15 000 ×*g* for 30 min. The supernatant was applied to Ni-NTA resin (Sigma) and washed with 10 column volumes of lysis buffer before eluting with 20 mM TrisHCl, 50 mM NaCl, 300 mM imidazole, 10% glycerol, 10 mM β-mercaptoethanol, pH 8. The IMAC pool was diluted 1:4 in 50 mM Tris HCl pH 8 and applied to 4 mL DEAE-Sepharose (Sigma). After washing with 20 mM Tris HCl, 50 mM NaCl, pH 8, proteins were eluted with a stepwise gradient of NaCl in 100 mM increments. Purity of the fractions was assessed by SDS-PAGE and protein concentration measured using a BCA assay (Pierce).

MalE-EndoN was expressed and purified as previously described (53).

### Sample preparation

Cells were resuspended in RIPA buffer (50 mM TrisHCl pH 7.5, 150 mM NaCl, 0.5 % Triton X-100, 0.5 % sodium deoxycholate, 0.1 % SDS) supplemented with 1 mM PMSF and 500 U benzonase and lysed at 4 °C for 30 min. Cell debris was pelleted at 18 000 ×*g* for 15 min at 4°C. The protein concentration of the supernatant was measured using a BCA assay (Pierce) and supernatants were to appropriate concentrations with RIPA buffer. For endosialidase assays, MalE-EndoN was added to a final concentration of 10 μg/mL and both treated and untreated samples were incubated at 37 °C overnight (ELISA). For blotting Mal-E-EndoN was added to a final concentration of 0.5 mg/mL and both treated and untreated samples were incubated at 37 °C for 30 min.

For determination of polySia glycan linkages, MCF-7 cells at ~ 40% confluency were treated with the O-linked glycosylation inhibitor, 2 mM benzyl 2-acetamido-2-deoxy-α-D-galactopyranoside (benzyl-GalNAc; Sigma), for three days. Supernatants from cells +/− benzyl-GalNAc were prepared as described above. 1200 μg protein was mixed with 15 μL Glycoprotein Denaturing Buffer (NEB) in 150 μL total volume and incubated at room temperature for 10 min. 30 μL 10 % NP-40, 30 μL Glyco Buffer 2, 90 μL dH_2_O and +/− 2500 U PNGase F (NEB) was added to each tube and incubated at 37 °C for 1.5 h. Samples were analysed directly by ELISA.

For secretion of polySia, MCF-7 cells at ~ 40% confluency were washed with PBS, then incubated with Opti-MEM (Gibco) supplemented with 30 mM glucose, 2 mM CaCl_2_, 40 mM non-essential amino acids (Thermo Fisher), 10 ng/mL hydrocortisone, and 1X penicillin-streptomycin at 37 °C for three days. Media was collected and floating cells were removed by centrifugation at 300 ×*g* for 10 min. The supernatant was spun at 2000 ×*g* for 10 min. At this point, the media was concentrated 10-fold (using Amicon 3 kDa molecular weight cutoff filters) for CD63 ELISAs or carried forward for isolation of extracellular vesicles. EVs were pelleted by ultracentrifugation at 100 000 ×*g* for 70 min at 2 °C. The EV-depleted media (supernatant) was retained and concentrated for analysis. The pellet (EVs) were resuspended in PBS and spun at 100 000 ×*g* for 70 min at 2 °C, then resuspended in 100 μL PBS.(54)

### Immobilization of GFP-EndoN_DM_ and isolation of polysialylated proteins

Freshly purified GFP-EndoN_DM_ (above) was dialyzed against 0.1 M NaHCO_3_, 0.5 M NaCl at 4 °C overnight then concentrated to 20 mg/mL using Amicon 50 kDa MWCO filters. The concentrated protein was immobilized on cyanogen bromide activated Sepharose 4G (Sigma) according to manufacturer’s instructions. The resin was stored in 0.1 M Tris HCl pH 8, 0.5 M NaCl at 4 °C.

MCF-7 cells were incubated with 10 μM Ac_4_ManNAz (Carbosynth) for 3 days to label sialic acids with azides. Cells were resuspended in RIPA buffer supplemented with 1 mM PMSF and 500 U benzonase and lysed at 4 °C for 30 min. Cell debris was pelleted at 18 000 ×*g* for 15 min at 4°C and the protein concentration of the supernatant was measured using a BCA assay (Pierce). Polysialylated proteins were enriched using 500 μL GFP-EndoN_DM_-conjugated Sepharose 4G resin and 1 mg total protein in 15 mL 25 mM TrisHCl pH 8, 600 mM NaCl at 4 °C overnight with gentle rotation. Polysialylated proteins were eluted by incubating twice with 200 μL 25 mg/mL colominic acid (Carbosynth) at room temperature for 1 – 2 h and collecting the supernatant. The resin was washed with a further 500 μL 25 mM Tris HCl pH 8, 600 mM NaCl and then the azido-sialic acids were labeled with 30 μM WS DBCO-biotin (Click Chemistry Tools) at room temperature for 30 min. Excess WS DBCO-biotin was removed through multiple rounds of filter centrifugation using a 3000 MWCO Amicon filter. Biotinylated proteins were captured using 200 μL streptavidin-agarose at 4 °C overnight with gentle rotation. The resin was washed extensively with 1 % SDS in PBS, then PBS to remove the detergent. Proteomic analysis was performed directly on the beads.

### Sample preparation and mass spectrometry

Samples were treated with PNGase F (Promega) according to manufacturer’s instructions, then dried. Samples were resuspended in 50 μL of 50 mM NH_4_HCO_3_ (pH=8.3), and cysteines were reduced with 10 mM DTT at 60°C for 1 hour. Samples were cooled to room temperature and iodoacetamide was added to a final volume of 20 mM. Samples were incubated at room temperature in the dark for 30 minutes. Iodoacetamide was then inactivated by adding DTT to a final concentration of 40 mM. MS grade TPCK-treated trypsin (Promega) was added to a final protease:protein ration of 1:50-1:100 and samples were digested overnight at 37 °C. Supernatant was removed from beads, lyophilized and re-suspended in 1% TFA. Peptides were purified by homemade C18 tips, and then lyophilized.

Samples were analyzed on a linear ion trap-Orbitrap hybrid analyzer (LTQ Velos-Orbitrap Elite, ThermoFisher, San Jose, CA) outfitted with a nanospray source and EASY-nLC split-free nano-LC system (ThermoFisher, San Jose, CA). Lyophilized peptide mixtures were dissolved in 0.1% formic acid and loaded onto a 75 μm × 50 cm PepMax RSLC EASY-Spray column filled with 2 μM C18 beads (ThermoFisher San, Jose CA) at a pressure of 800 BAR. Peptides were eluted over 120 min at a rate of 250 nL/min using a gradient set up as 0 % – 30 % gradient of Buffer A (0.1 % Formic acid; and Buffer B, 0.1 % Formic Acid in 100 % acetonitrile). Peptides were introduced by nano electrospray into an LTQ-Orbitrap hybrid mass spectrometer (Thermo-Fisher). The instrument method consisted of one MS full scan (400–1500 m/z) in the Orbitrap mass analyzer, an automatic gain control target of 1e6 with a maximum ion injection of 200 ms, one microscan, and a resolution of 240,000. Ten data-dependent MS/MS scans were performed in the linear ion trap using the ten most intense ions at 35% normalized collision energy. The MS and MS/MS scans were obtained in parallel fashion. In MS/MS mode automatic gain control targets were 3e5 with a maximum ion injection time of 50 ms. A minimum ion intensity of 5000 was required to trigger an MS/MS spectrum. The dynamic exclusion was applied using a maximum exclusion list of 500 with one repeat count with a repeat duration of 30 s and exclusion duration of 20 s. Peptides were identified using a human peptide atlas and data was visualized using Scaffold v4.

### ELISA

GFP-EndoN_DM_ was allowed to adhere to 96 well Nunc-Immuno plates (ThermoFisher) in 100 μL at 100 μg/mL protein diluted in PBS at 4 °C overnight in a humidified container. The overnight incubation is simply to make the following day shorter and can be decreased to 1 h at room temperature. All subsequent incubations were performed at room temperature for 1 – 2 h and all washing steps were performed three times with 1X PBS, 0.5 % tween-20 (PBS-T). Plates were treated with the following solutions with washes between each: 300 μL 2.5 % BSA in PBS-T to block, 0.01 – 100 μg/mL colominic acid or 0.4 – 6 mg/mL protein from serum, lysates or reactions in 100 μL, 100 μL primary antibody, and 100 μL secondary antibody. The amount of protein per well and dilutions of primary antibodies were 40 μg and 1/1000 for QSOX2, 100 μg and 1/10 000 for GOLIM4, 600 μg and 1/100 for AGR2, 600 μg and 1/100 for G6PD, 600 μg and 1/100 for MGAT2, 600 μg and 1/100 for LRP2, and 600 μg and 1/100 for SDF4. The CD63 primary antibody was used at 1/500 After a final three washes, 100 μL 3,3' 5,5'-Tetramethylbenzidine (TMB; ThermoFisher) was added to each well and absorbance was measured at 652 nm every 5 min. Error bars denote standard deviations on biological replicates run at least three times in technical duplicate.

### Microscopy

MCF-7 cells grown on coverslips to ~90 % confluency. All washing steps were performed three times with PBS. Cells were fixed with ice cold 2% formaldehyde in MeOH at −20 °C for 5 min (PMID: 10098197). After washing, coverslips were blocked in 3 % BSA, PBS-T at room temperature for 1 h, then incubated with either 1:1600 α-GM130 or 1:100 α-QSOX2 (Abcam ab121376) in 3 % BSA, PBS-T at 4 °C overnight. Coverslips were washed and then incubated with 1:500 AlexaFluor 633 goat α-rabbit IgG (A21070, Thermo Fisher Scientific), 0.1 mg/mL GFP-EndoN_DM_ at room temperature in the dark for 90 min. Coverslips were washed and then labeled with ActinRed 555 (Invitrogen) and NucBlue (Invitrogen) according to manufacturer’s instructions before inverting into a drop of CytoSeal 60 (Thermo Fisher Scientific). Analysis was performed using a Zeiss spinning disk confocal microscope.

### Western blotting

For each sample, equivalent amounts of protein were loaded on SDS-PAGE gels and transferred to PVDF membrane. Membranes were blocked in 5% BSA, 1X TBS-tween, then incubated with primary antibody in blocking buffer at room temperature or 4 ℃ overnight. Dilutions of primary antibodies were 1/1000 for polySia, 1/500 for QSOX2, 1/10 000 for GOLIM4, 1/500 for AGR2, 1/4000 for β-tubulin. After washing in 1X TBS-tween, membranes were incubated with appropriate 1:1000 anti-mouse IgG-HRP or 1:2000 anti-rabbit IgG-HRP for 1 h, then washed and developed using Pierce ECL western blotting substrate.

### Flow Cytometry

10 μl EVs were stained with several different concentrations of GFP-EndoN_DM_ at room temperature in the dark for 90 min. Separate control tubes were prepared for each GFP-EndoN_DM_ concentration. Control samples containing PBS only or unstained EVs were also prepared. EV samples and controls were diluted with 0.22 μm filtered PBS and a dilution series of each was prepared. Samples were acquired on a Cytoflex LS cytometer using a violet-SSC configuration. The instrument was calibrated before each run using a mixture of bead sizes (Apogee) to ensure appropriate settings and separation of discrete sized particles. Each sample condition was compared to the appropriately diluted unstained EVs and GFP-EndoN_DM_ only controls to clearly identify non-specific aggregates. Analysis was performed in FlowJo v10.

## Results

### Polysialic acid is secreted from the MCF-7 breast cancer cell line

MCF-7 cells have long been known to contain intracellular polySia (55) but the context was not defined. We confirmed that polySia found in MCF-7 cells is intracellular and localized predominantly in the Golgi apparatus, an integral component of the secretory pathway (Fig. 1A). To determine whether polySia was secreted, MCF-7 cells were washed with PBS to remove any interference from polySia in fetal bovine serum and then incubated in serum free Opti-MEM. The concentration of polySia in the culture media was significantly increased after three days incubation (Fig. 1B), as determined by ELISA with polySia-specific lectin EndoN_DM_ as the capture reagent and polySia antibody 735 for detection (42). Western blotting of the ER-resident control protein disulfide isomerase (PDI) demonstrated that the polySia in culture media was not a result of lysed cells (Fig. 1C).

**Figure 1:**
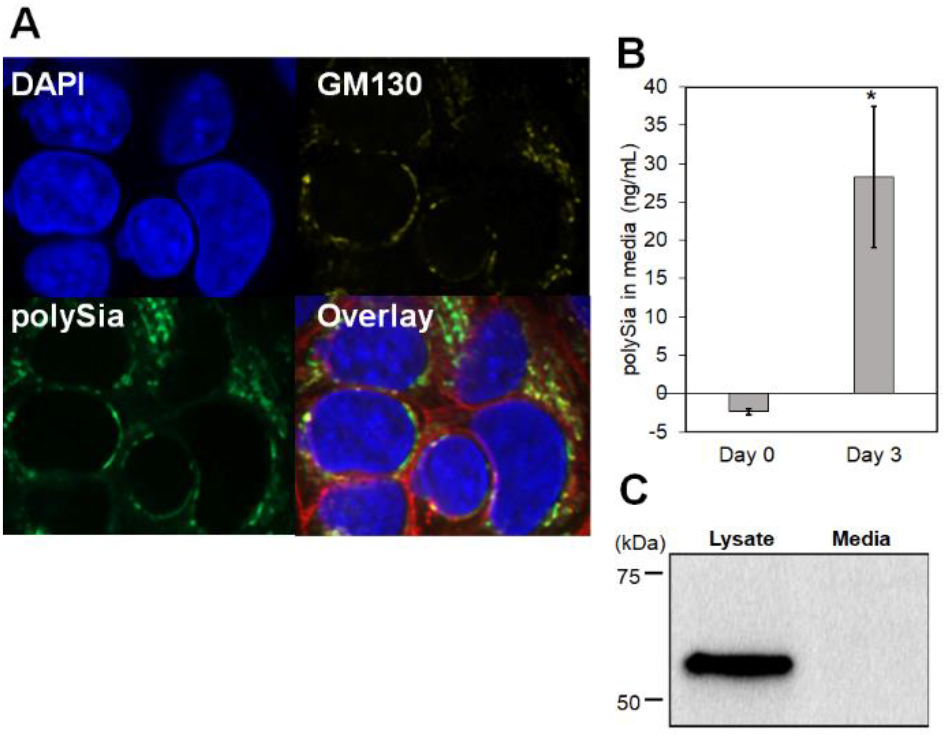
MCF-7 cells secrete intracellular polysialic acid. A) Confocal microscopy of MCF-7 cells. PolySia (green) is intracellular and is co-localized with Golgi marker GM130 (yellow). Nucleus is stained with DAPI (blue) and red in overlay is actin. B) polySia ELISA of MCF-7-conditioned Opti-MEM showing secretion of polySia into media. C) α-PDI immunoblot of cell lysate and cell media. Equivalent amounts of protein are loaded in both wells. Absence of PDI indicates that the polySia in the media cannot be attributed to dead cells.

The secreted polySia observed could be a component of extracellular vesicles (EVs) or a soluble glycoprotein. To characterize the secreted polySia we isolated EVs from cell conditioned Opti-MEM by ultracentrifugation and retained the EV-depleted culture media. Immunoblots comparing the isolated EVs with the EV-depleted media identified polySia on both secreted fractions, with most of the polySia in the soluble (EV-depleted) fraction (Fig 2A). Treatment with the polySia-specific hydrolase EndoN abolished the signal, confirming the specificity of the immunoblotting (Fig 2A). The small fraction of EV-associated polySia could not be observed on comparing the conditioned media to the conditioned media depleted of EVs (Fig 2B). To support the observation that some of the secreted polySia is associated with EVs, we examined the conditioned Opti-MEM using a variation of the polySia ELISA, where after the adsorbed polySia lectin (EndoN_DM_) captures polysialylated species, including EVs, they can then be detected with an antibody specific for the EV marker CD63 (56–58) (Fig S1). Analysis of the conditioned Opti-MEM showed that polySia-associated CD63 increased over time (Fig S2), demonstrating that this variation of the polySia ELISA can detect polysialylated EVs secreted from cells. Tosupport the observation that polySia is secreted from MCF-7 cells both on EVs and on soluble proteins, we measured the amount of polySia and polySia-associated CD63 in the media after ultracentrifugation to pellet EVs (Fig. 2B). While we observed the expected decrease in polySia-associated CD63 after ultracentrifugation, we did not detect a change in total polySia (Fig. 2B), consistent with out polySia immunoblot suggesting that most of the polySia secreted from MCF-7 cells is attached to soluble proteins (Fig. 2A). To validate our observation that polySia occurs on EV surfaces, as opposed to insoluble protein, we analyzed isolated EVs directly by flow cytometry and observed polySia signal in the EV species (Fig 2C). This multi-technique approach to characterizing polySia in the MCF-7 cell secretome provides strong evidence of polySia on both soluble and EV-associated glycoproteins.

**Figure 2:**
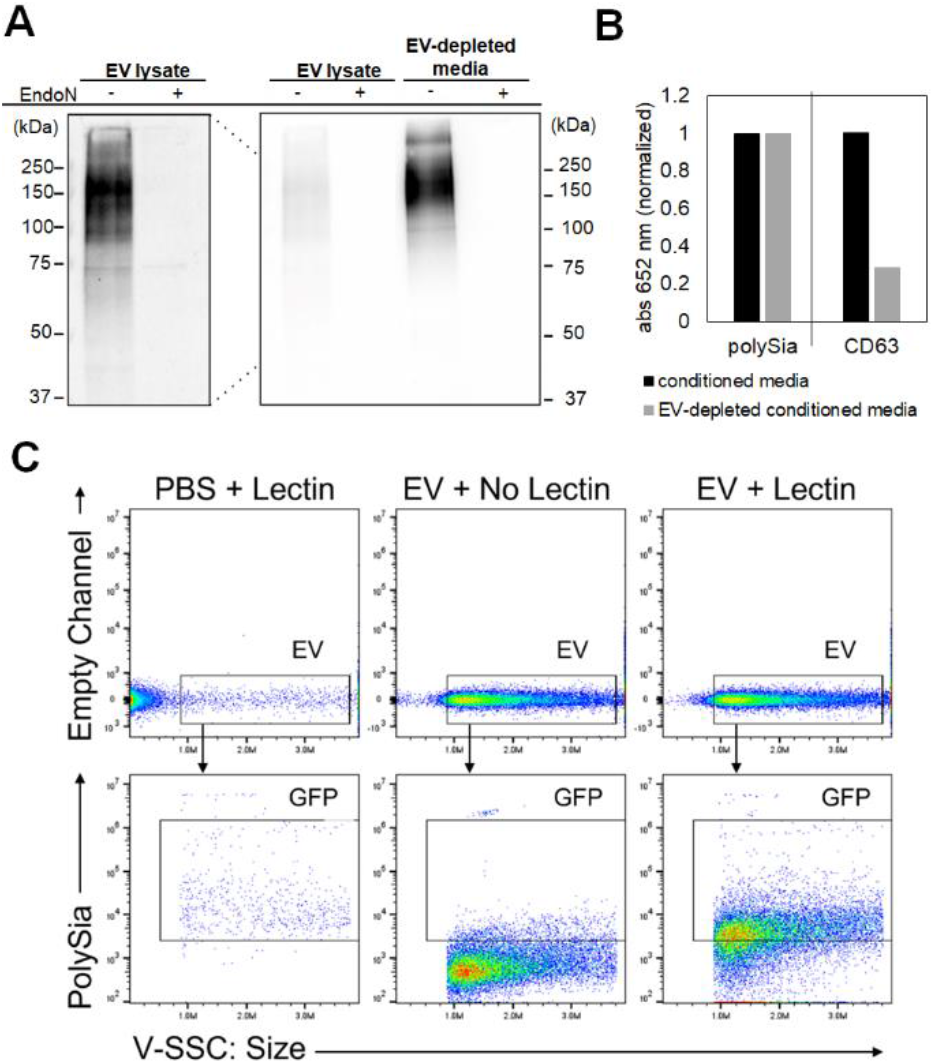
Polysialic acid is secreted from MCF-7 cells on extracellular vesicles and soluble proteins A) polySia immunoblot of secreted polySia. EVs were isolated from conditioned Opti-MEM and the EV-depleted media was retained and concentrated. Protein concentration was normalized using the BCA assay and an equivalent amount of protein was loaded into each well. The panel on the left is an overexposure of the panel on the right. B) ELISA comparing MCF7 conditioned Opti-MEM before and after EV depletion. C) Flow cytometry of EVs isolated from MCF-7 cells stained with fluorescent polySia-specific lectin GFP-EndoN_DM_, including appropriate lectin only and EV only controls.

### Identification and validation of polysialylated proteins in MCF-7 cells

To improve the isolation of polysialylated proteins, we developed a two-stage protocol comprised of an affinity purification step and introduction of a bioorthogonal handle for covalent modification and subsequent isolation (Fig 3A). First, the bioorthogonal handle was introduced by growing MCF-7 cells in the presence of Ac_4_ManNAz, a modified precursor of sialic acid which is metabolically incorporated into sialylated and polysialylated glycoproteins (59, 60). The concentration of Ac_4_ManNAz was optimized such that it was sufficient to ensure good incorporation but not so high that it reduced the affinity of polySia for the EndoN_DM_ lectin (Fig S3). Different proteins may differentially incorporate 5-azido-sialic acid however, which could lead to different affinities for the lectin. The polysialylated proteins were next separated from other sialylated proteins and bulk cellular components by affinity purification using immobilized GFP-EndoN_DM_ and eluting with free polySia (colominic acid). Lastly, the bioorthogonal handle was used to separate polysialylated proteins from non-specifically bound proteins by condensing the eluate with a biotin-cyclooctyne conjugate which installs biotin at azide-bearing sialic acids. Biotinylated proteins were then isolated on streptavidin-agarose and subjected to high-stringency washes in 1 % SDS. Proteomic analysis of peptides released from streptavidin beads following PNGase F treatment to remove *N*-linked glycans yielded 31 proteins with two or more exclusive peptides (Table S1). These proteins were examined against the Contaminant Repository for Affinity Purification (CRAPome) database (61) to filter out those that interact non-specifically with agarose beads (Table S1). Seven proteins were identified as being present in 5% or less of the CRAPome experiments: QSOX2, GOLIM4, AGR2, G6PD, MGAT2, SDF4, and LRP2 (megalin). To validate the presence of polySia on the identified proteins, we analyzed MCF-7 cell lysate using the polySia ELISA with antibodies specific for the protein of interest as the detection antibodies (Fig 3B). Notably, the cells were lysed using RIPA buffer to disrupt the cell membranes, so signal indicates that the proteins themselves are polysialylated. Three of the proteins, QSOX2, GOLIM4, and AGR2, gave robust signal which was substantially reduced upon treatment with the polySia-specific hydrolase EndoN (Fig 3B). For the remaining four proteins, signal was much lower but did also decrease when treated with EndoN, making it more difficult to unambiguously determine whether they are polysialylated.

**Figure 3:**
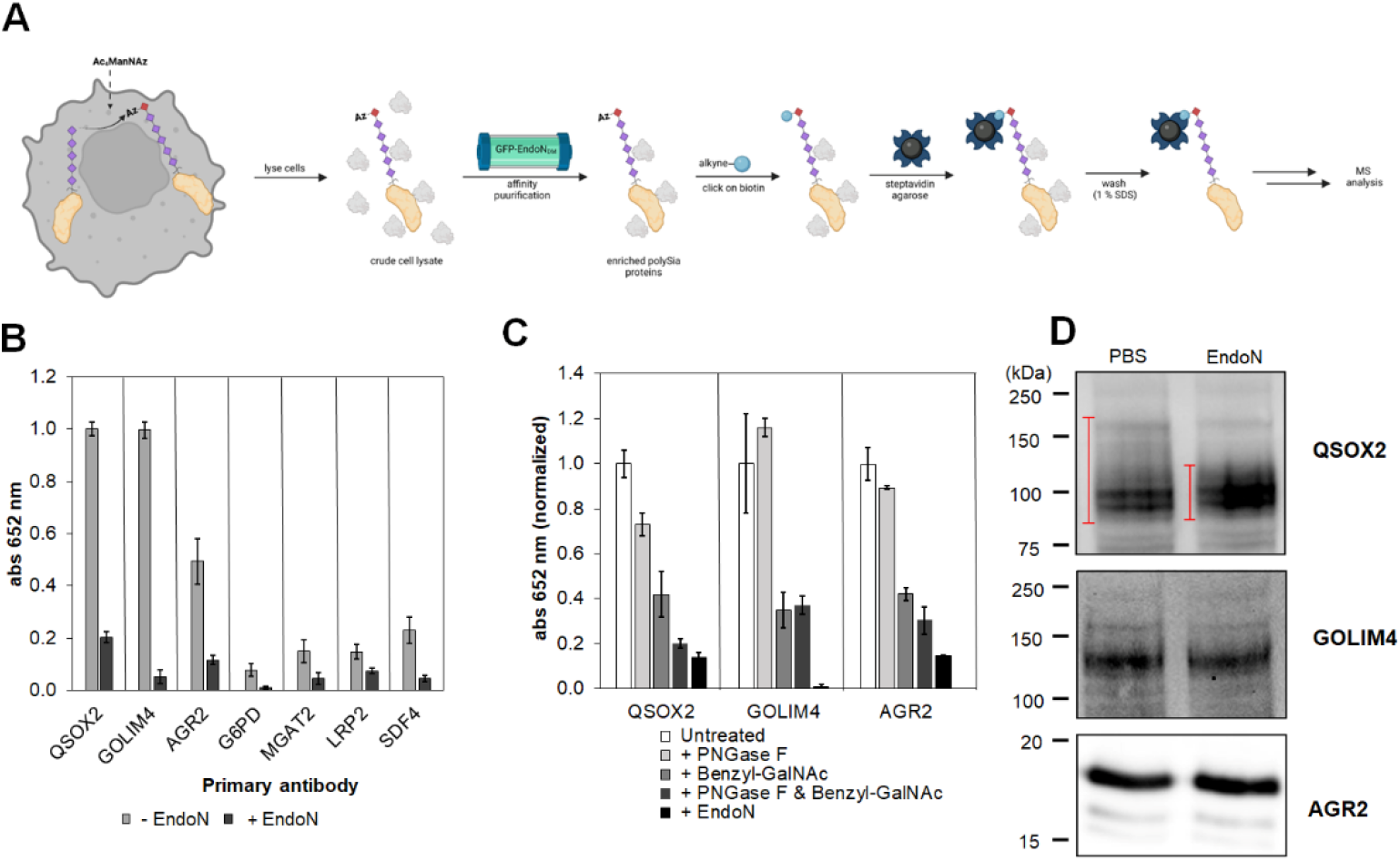
Improved methodology to identify polysialyalated proteins. A) Workflow for isolating polySia proteins. Made with www.Biorender.com B) ELISA validation of polysialylated proteins identified in MCF-7 cells. ELISA plates were coated with polySia lectin GFP-EndoN_DM_ to capture polySia in MCF-7 cell lysate. PolySia was removed by treatment with EndoN prior to analysis. B) Identification of *N*- vs *O*-linked polySia in validated polySia proteins. C) ELISA determining whether polySia is attached to N-linked glycans (enzymatically removed with PNGase F) and/or O-linked glycans (metabolically inhibited with benzyl-GalNAc). D) Immunoblots of validated polySia proteins in MCF-7 cell lysate. Red bars show spectrum of polysialylated QSOX2.

The linkage of glycans carrying polySia was determined for QSOX2, GOLIM4 and AGR2 proteins using the O-linked glycosylation inhibitor, benzyl-GalNAc, and the N-glycosidase, PNGase F (62, 63). Cells were incubated for two days with or without benzyl-GalNAc to inhibit formation of *O*-linked glycans, then lysates from these cells were treated with PNGase F for 1 h after denaturation to remove *N*-linked glycans. Both benzyl-GalNAc and PNGase F caused a decrease in signal for polySia-QSOX2 detected by polySia-ELISA and the combination of the two treatments is equivalent to treatment with EndoN, suggesting that both N- and O-linked glycans on QSOX2 are polysialylated (Fig 3C). For polySia-GOLIM4 and -AGR2, signal was decreased upon treatment with benzyl-GalNAc only, suggesting polySia occurs on O-linked glycans in these proteins (Fig 3C).

To compliment the ELISA, the mobility of QSOX2, GOLIM4, and AGR2 on western blot was determined before and after treatment of the cell lysates with EndoN to remove polySia. The expectation for highly polysialylated proteins, such as NCAM, is that the glycoprotein would be a high molecular weight smear which would collapse to a band closer to the expected molecular weight of the protein after treatment with EndoN (Fig. S4). However, NCAM has two N-linked tetra-antennary glycans meaning each protein can conceivably carry up to eight polySia chains. Proteins with fewer polySia chains, partial polysialylation and/or short polySia chains may not yield gel shifts as substantial as those observed for NCAM. QSOX2 exhibited a clear decrease in the apparent molecular weight after treatment with EndoN, (Fig. 3D, S5), supporting the ELISA data that it is polysialylated. Neither GOLIM4 nor AGR2 showed a shift in apparent molecular weight upon treatment with EndoN (Fig. 3D), despite clear results from the ELISA that both can be polysialylated. The ambiguity of the immunoblots for AGR2 and GOLIM4 could be due to modification of only a small fraction of the protein with polySia, making the high molecular weight smear from the polysialylated fraction indistinguishable from the background. The polysialylation of only a small fraction of a protein was previously observed with NRP2 and ESL-1 in mouse microglia and human macrophages (41). In contrast to immunoblotting our ELISA method only detects proteins that are polysialylated, making it a more unequivocal tool for the characterization of polySia proteins. The milder conditions of the ELISA compared to SDS-PAGE and immunoblotting are advantageous since some tertiary or even quaternary structures can be maintained and polySia doesn’t necessarily interfere with detection.

### MCF7 cells secrete polysialylated QSOX2

With the identification of three novel polysialylated proteins we investigated whether these proteins were secreted from MCF-7 cells in soluble form and as components of EVs. Because of the limited abundance of secreted proteins, particularly in isolated EVs, we were restricted to using immunoblotting rather than the polySia ELISA. We did not observe any staining for GOLIM4 in either secreted fraction (data not shown). AGR2 was identified in the EV lysate but not in the EV-depleted media (Fig 4). The anti-AGR2 immunoblots did not reveal whether secreted AGR2 was polysialylated, likely for similar reasons to those discussed above. In contrast, QSOX2 was identified in both the EV lysate and the EV-depleted media (Fig 4). Treatment of the EV-depleted media with EndoN resulted in the collapse of the smeared signal to a lower molecular weight band, confirming that secreted QSOX2 is also polysialylated.

**Figure 4:**
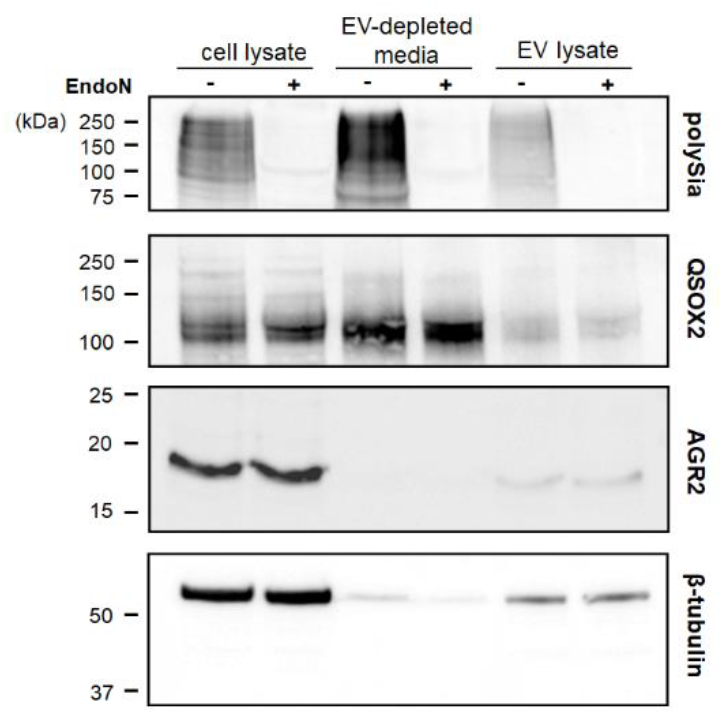
Immunoblot analysis of MCF-7 secretome. Protein concentration of MCF-cell lysate, EV-depleted media, and EV lysate were determined using the BCA assay and an equivalent amount of protein was added in each well.

## Discussion

PolySia expression has a well-established correlation with cancer severity, but the mechanistic basis for this effect has not been elucidated. Recent reports that detected polySia in intracellular pools and in cell culture media from heathy cells (40, 41) led us to hypothesize that polySia may also be secreted from cancer cells. Herein we provide evidence that polySia is secreted from MCF-7 breast cancer cells. We identified polySia in both EV-depleted media and on EVs which suggested a multifaceted role for secreted polySia. Based on RNA expression data from the Human Protein Atlas (www.proteinatlas.org) (64) the MCF-7 cell line does not express any of the proteins known to be polysialylated. Therefore, to gain biological context for this secreted polySia we searched for novel polysialylated proteins.

Identification of polysialylated proteins is challenging because of the large size and anionic charge on the polymer. Many proteins, particularly basic proteins, tend to interact non-specifically with polySia and cannot be removed under the mild washing conditions of immunoprecipitation. The resulting proteomics experiments often contain numerous hits which require either a substantial number of antibodies for validation or a biased selection of hits to validate based on known function and subcellular localization. Previous attempts to identify polysialylated proteins relied on immunoprecipitation experiments with the α-polySia mAb (10, 65). The resulting hits were then sorted manually for proteins that are likely to be on the cell surface, where polySia is typically localized. This biased approach leads to the discarding of proteins which may have more than one function. For example, MGAT2, which we have tentatively identified as polysialylated in MCF-7 cells, has previously shown up in α-polySia proteomic experiments but was discarded because it is not a cell surface protein (10).

The addition of Ac_4_ManNAz, which introduced bioorthogonal functionality into polySia, allowed for stringent washing steps after labeling with biotin/streptavidin. Our approach to finding polysialylated proteins is similar to one used for identification of proteins carrying the post-translational modification *O*-GlcNAc. Cells were incubated with an azido-modified precursor, Ac_4_GlcNAz, and the resulting proteomic analysis identified over 100 putative O-GlcNAcylated proteins, 23 of which were confirmed by reciprocal immunoprecipitation (66, 67). Compared to proteins that carry *O*-GlcNAc, a carbohydrate moiety composed of a single monosaccharide, proteomics of polysialylated proteins is further complicated by the fact that polysialylated proteins are frequently highly glycosylated, and highly glycosylated proteins are generally incompatible with mass spectrometry methods for identifying proteins. Glycans reduce the efficiency of the trypsin digest, the number of peptides observed after mass spectrometry, and the ability of the software to accurately identify a peptide (68). While N-linked glycans can be removed before analysis, the same is not true for *O*-linked glycans. The benefit of our methodology to enrich the polysialylated proteins, combined with stringent washing and CRAPome filtering, is evidenced by the fact that out of the seven hits, three were fully validated. The remaining four hits were all positive for polySia according to our ELISA, but the signal was not robust enough for unambiguous validation. No other single study has identified more than one new polysialylated protein, which could reflect the increased efficiency of this method, though this could also be representative of the fact that cancer cells express more polysialylated proteins than healthy cells.

All of the polysialylated proteins identified in this study can be membrane associated, as would be predicted for interaction with polysialyltransferase enzymes. All have also been detected in human serum, indicating they can be secreted in healthy humans (69–71). GOLIM4 is a Mn^2+^-binding protein that cycles between endosomes and the Golgi (72), which also acts as the receptor for Shiga toxin trafficking to the Golgi (73). Along with NCAM and ST8SiaIV, two other known polysialylated proteins, GOLIM4 is a target for the β-site APP cleaving enzyme 1 (BACE1) secretase, the aspartyl protease that is responsible for cleavage of amyloid precursor protein in Alzheimer’s disease (74, 75). The function of the BACE1 secretase outside of APP cleavage is not known, but the lower 60 kDa band observed in α-GOLIM4 western blot (Fig S6) could be a product of this proteolysis. It is also possible that there are proteolytic fragments not recognized by the antibodies used in this study. We did not observe GOLIM4 secretion from MCF-7 cells, despite its previous identification in microparticles (70). Cells growing in a petri dish may not experience the same pressures to secrete specific proteins. Consequently, while we did not observe secretion of GOLIM4 into media, it is possible that it may be secreted under certain conditions or by different cell types.

We did observe secretion of the other two identified polySia-proteins from the MCF-7 cells. Both QSOX2 and AGR2 immunostaining was observed in isolated EVs, and robust QSOX2 immunostaining was observed in EV-depleted conditioned Opti-MEM. AGR2 is a disulphide bond isomerase that is strongly upregulated in many cancers, where it correlates with metastasis (71, 76). It is normally ER resident as it has a KTEL retention sequence but can also be *O*-glycosylated and secreted (71, 77). The results of our linkage analysis suggests that polySia is primarily on *O*-glycans on AGR2. The presence or absence of polySia on secreted EV-associated AGR2 was not determined in this study due to sample size limitations. QSOX2 is a sulfhydryl oxidase which makes disulfides de novo. It is a relatively unstudied protein but is beginning to gain attention in the context of cancer. In a genetic study of colorectal cancer, QSOX2 overexpression was a predictor of poor prognosis (78). Knockdown of QSOX2 significantly reduced proliferation of colorectal cancer (79) and non-small cell lung cancer cells (80), and it was also identified as a potential serum biomarker for lung cancer (80). QSOX2 also shares 50 % similarity with QSOX1, which is more studied and has recently been identified as a marker of metastasis in several cancers, including prostate (81, 82), pancreatic (36, 83), and breast (84, 85) cancers. QSOX1 localizes to the ER and Golgi but can be secreted (83, 86–88), in a mechanism that requires its N-linked glycan (89). Here we have demonstrated that QSOX2 carries polySia on both *N*- and *O*-linked glycans, and that secreted QSOX2 is polysialylated.

The role of polySia in the function of these proteins in the context of cancer biology and the cancer secretome remains in question. PolySia has multiple potential mechanisms by which it promotes cancer progression and metastasis. It may be that polySia enhances serum half-life of these proteins (90) or causes their targeting to the metastatic niche. It is possible that polySia may dampen the immune response in patients with cancer. PolySia could also sequester proinflammatory cytokines like IL-6, which also promote metastasis (91). More research is needed to understand the role of polySia in the function of these proteins. From a diagnostic biomarker perspective, the polysialylation of these proteins introduces exciting opportunities. Recent analysis of cancer secretomes suggest that they contain unconventional proteins (92) which make for good candidates for diagnostic biomarkers because these proteins would not otherwise be found in healthy serum. Because polySia has restricted expression in healthy adults and aberrant expression in cancer, it is possible that proteins like QSOX2 are not polysialylated in healthy individuals. Detection of the polysialylated fraction of these secreted proteins from cancer cells may improve the diagnostic or prognostic potential of these proteins.

### Contribution to the Field Statement

The glycan polysialic acid plays integral roles in health and disease. However, the mechanisms by which it controls biology are poorly understood, due in large part to a lack of analytical tools with which to study it. We developed new methodology to identify polysialylated proteins which overcomes the limitations of immunoprecipitation. Using this methodology we identified three new polysialylated proteins in the MCF-7 cell line alone, adding to the current list of 5-9. We demonstrated that polysialylated proteins can be secreted from cancer cells on their own and as part of extracellular vesicles. This observation, combined with two previous reports of polySia secretion from healthy cells, suggests that secretion of polySia may be a general phenomenon and bears further investigation. Two of the newly identified proteins are being investigated for their strong association with metastasis and poor prognosis in cancer patients. Given that polySia is also strongly associated with metastasis and poor prognosis, it is possible that polysialylation of these proteins contributes to their pathological activity and may even represent a potential diagnostic marker for cancer progression.

## Acknowledgements

This work was supported by grants to M.N. and L.W. from the Canadian Glycomics Network and M.N. from U.S. Department of Defense Breast Cancer Research Program, Breakthrough Award (#12045028), as well as a Banting Postdoctoral Fellowship and NSERC Discovery grant (RGPIN-2021-02888) awarded to L.W. We would like to thank Jonathan Krieger at SPARC Biocenter for advice on the proteomics analysis.

## Abbreviations

AGR2: anterior gradient protein 2
BACE: β-site APP cleaving enzyme 1
CCR7: C-C chemokine receptor type 7
DMB: 1,2-diamino-4,5-methylenedioxybenzene dihydrochloride
G6PD: glucose-6-phosphate dehydrogenase
GM130: Golgi marker 130
GOLIM4: Golgi integral membrane protein 4
HRP: horseradish peroxidase
LRP2: low-density lipoprotein receptor-related protein 2
MGAT4: UDP-N-acetylglucosamine: α-1,3-D-mannoside β-1,4-N-acetylglucosaminyltransferase IV
NCAM: neural cell adhesion molecule
PDI: protein disulfide isomerase
polySia: polysialic acid
QSOX2: quiescein sulfhydryl oxidase 2
SDF4: stromal derived factor 4

## Supplementary Material

### 1.1 Supplementary Figures

**Figure S1.**
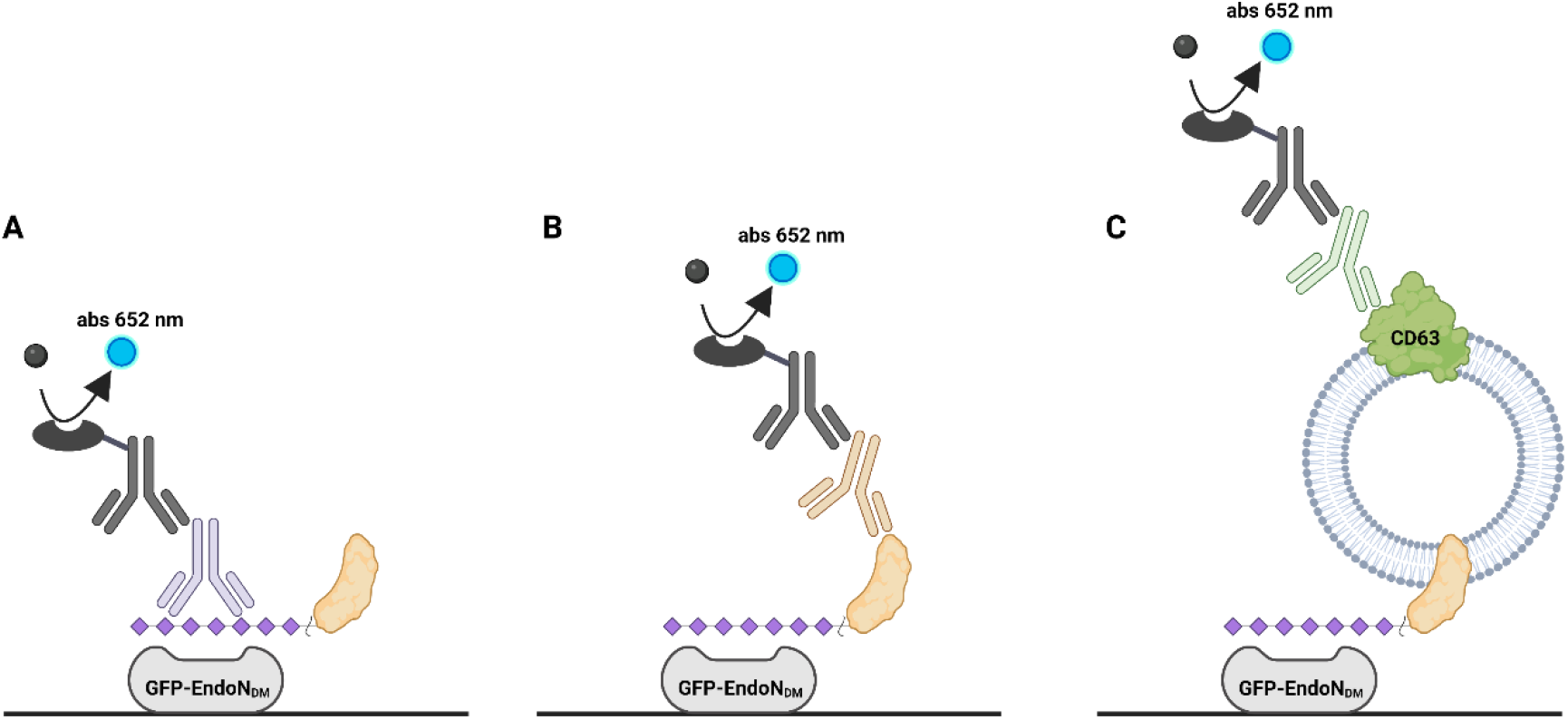
Schematic showing three possible versions of the polySia ELISA. Possible detections are polySia (A), a polysialylated protein (B), or polysialylated vesicles (C). Made with www.Biorender.com

**Figure S2.**
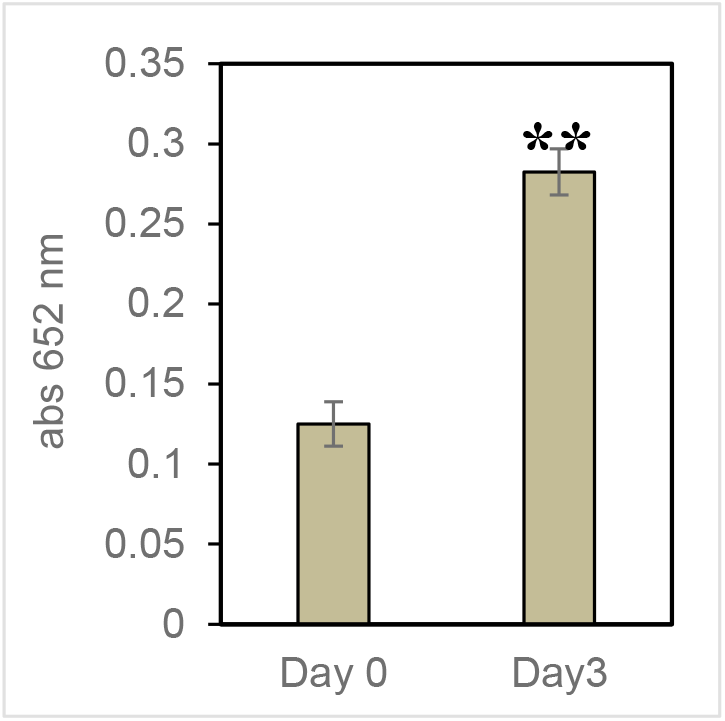
Polysialylated extracellular vesicles are secreted from MCF-7 cells. CD63-associated polySia was detected in conditioned Opti-MEM using the polySia ELISA.

**Figure S3.**
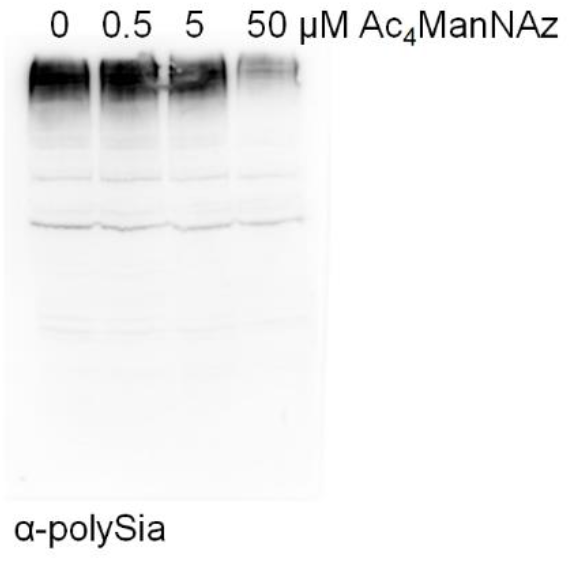
α-PolySia immunoblot showing that antibody reactivity is lost at concentrations of Ac_4_ManNAz higher than ~5 μM.

**Figure S4.**
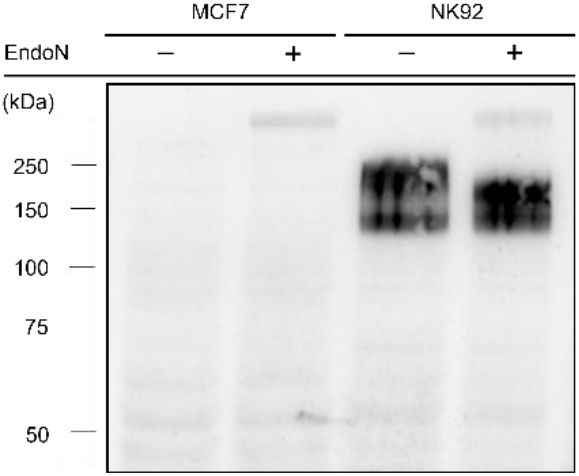
MCF-7 cells do not express NCAM. α-NCAM immunoblot of lysates from MCF-7 and NK92 (positive control).

**Figure S5.**
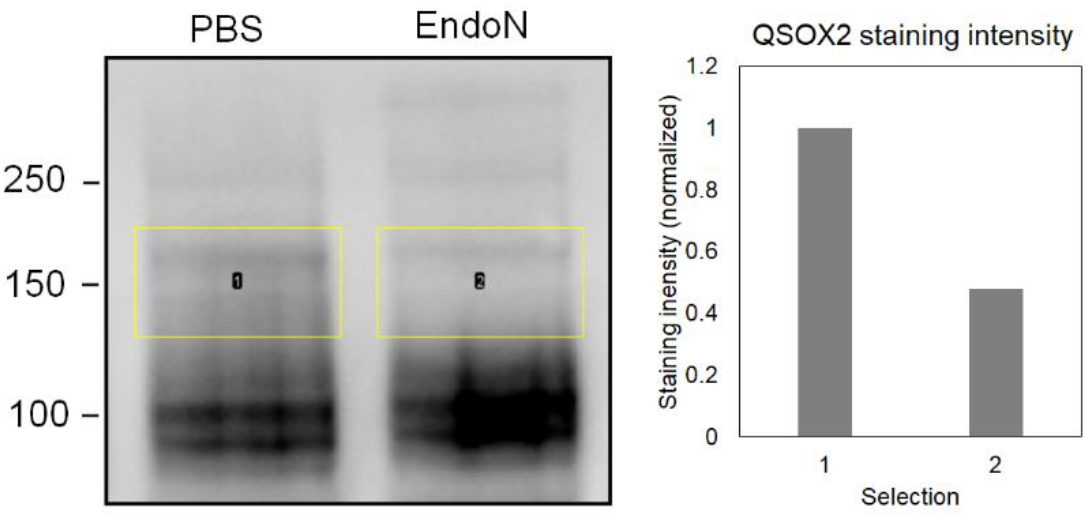
Densitometry analysis of QSOX2 immunoblot demonstrating QSOX2 is polysialylated. Densitometry was performed on the space within the yellow boxes. The resulting staining intensity, represented on the right as a bar graph, was normalized to the PBS-treated (polysialylated) samples.

**Figure S6.**
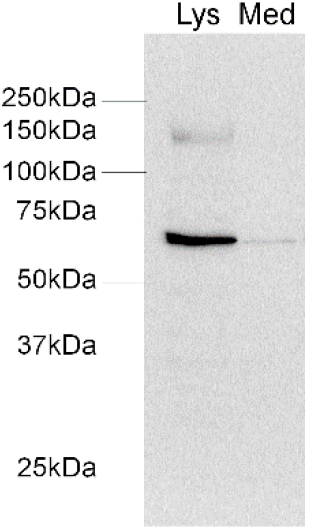
GOLIM4 may be secreted from MCF-7 cells. α-GOLIM4 immunoblot showing a potential proteolysis fragment that is secreted into Opti-MEM.

## Bio RENDER

49 Spadina Ave. Suite 200

Toronto ON M5V 2J1 Canada

www.biorender.com

### Confirmation of Publication and Licensing Rights

**July 19th, 2022**

**Science Suite Inc.**

***Subscription:** Individual*

***Agreement number:** QM246G0BKN*

***Journal name:** Frontiers in Molecular Biosciences*

To whom this may concern,

This document is to confirm that Lisa Willis has been granted a license to use the BioRender content, including icons, templates and other original artwork, appearing in the attached completed graphic pursuant to BioRender’s [Academic License Terms](https://biorender.com/academic-license/). This license permits BioRender content to be sublicensed for use in journal publications.

All rights and ownership of BioRender content are reserved by BioRender. All completed graphics must be accompanied by the following citation: “Created with BioRender.com”.

BioRender content included in the completed graphic is not licensed for any commercial uses beyond publication in a journal. For any commercial use of this figure, users may, if allowed, recreate it in BioRender under an Industry BioRender Plan.

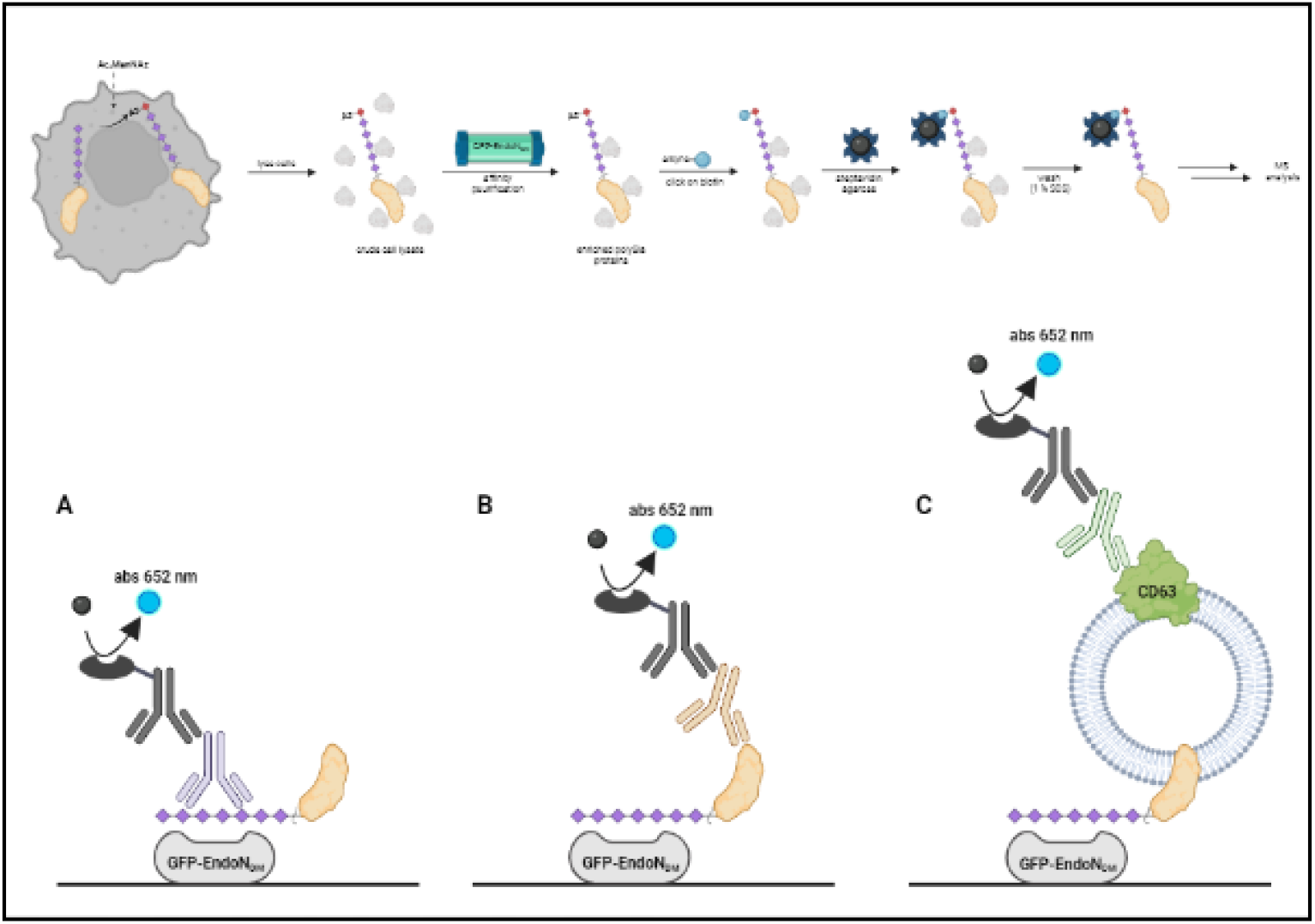

*For any questions regarding this document, or other questions about publishing with BioRender refer to our [BioRender Publication Guide](https://help.biorender.com/en/articles/5256792-biorender-publication-guide), or contact BioRender Support at.*

